# Nanoscale Core–Rim Compartmentalization of Histamine and GABA within Individual Synaptic Vesicles

**DOI:** 10.64898/2026.08.25.747175

**Authors:** Kunio Fujiwara

## Abstract

Classical synaptic transmission assumes that amino acid neurotransmitters and monoamines are stored in distinct vesicle populations. Here, we provide structural evidence for a dual-transmitter organization in which histamine and GABA are spatially partitioned into a core and rim within individual synaptic vesicles. Using an engineered glutaraldehyde–NaBH₄ immuno-electron microscopy platform, quantitative analysis of 2,195 vesicles revealed a previously unrecognized nanoscale architecture, with histamine-associated signal concentrated within the vesicular core (95.1%), while GABA is organized toward the peripheral rim. Light microscopy further demonstrated extensive histamine–GABA correspondence across central and peripheral tissues, including sympathetic ganglia and adrenal medulla. This intravesicular segregation challenges the conventional separation of amino acid and monoamine storage into distinct vesicle classes and reveals that chemically distinct transmitters can occupy organized domains within a single vesicular lumen. This architecture may provide a structural basis for the distinct physiological modes of GABAergic and histaminergic signaling, linking nanoscale vesicular organization to their established differences in temporal action.

**One-sentence summary:** Using engineered glutaraldehyde-NaBH4-based ultrastructural analysis, we identified a novel "core histamine–rim GABA" vesicular architecture within GABAergic neurons, fundamentally redefining traditional models of dual-transmitter co-packaging and release dynamics.

## Introduction

For more than half a century, synaptic vesicles have been broadly divided into two canonical classes: clear vesicles containing amino acid transmitters that mediate rapid signaling^1^, and dense-core vesicles storing monoamines associated with slower neuromodulation^2^. This classification has implicitly assumed that chemically distinct neurotransmitters are segregated into distinct vesicle populations. Yet whether different neurotransmitters can be spatially organized within the lumen of a single synaptic vesicle has remained unknown. In particular, it is unclear whether a small synaptic vesicle can accommodate chemically distinct transmitter domains without membrane-bounded internal compartments.

Histamine has traditionally been regarded as a monoaminergic neuromodulator largely restricted to the tuberomammillary nucleus (TMN)^3^. Its distribution outside classical hypothalamic pathways has remained difficult to assess, in part because of technical limitations in the ultrastructural detection of this highly labile small molecule. Using an engineered glutaraldehyde–NaBH₄ epitope-preservation platform, we overcome these limitations and directly visualize histamine at the electron-microscopic level. This approach reveals a previously unrecognized core–rim architecture within individual synaptic vesicles, with histamine concentrated in the vesicular core and GABA associated with the surrounding peripheral rim. This organization challenges the long-standing segregation of amino-acid and monoamine transmitters into distinct vesicle classes and demonstrates that chemically distinct transmitters can occupy spatially organized domains within a single small synaptic vesicle. Strikingly, this core–rim architecture is observed across central inhibitory circuits and peripheral tissues, including adrenal chromaffin cells and sympathetic ganglia. The distinct physiological properties of GABAergic and histaminergic signaling have long been recognized, with GABA mediating rapid inhibition and histamine producing slower, sustained neuromodulatory effects. However, the basis for these distinct temporal properties has remained unresolved for decades. The core–rim organization identified here may provide an explanation for this long-standing physiological question.

## Results and Discussion

**Electron microscopy** identified a previously unrecognized polarized core–rim architecture within synaptic vesicles of conventional GABAergic neurons (**<u>Figure 1</u>**). This organization consists of a dense histamine core surrounded by the well-established peripheral GABA rim (**<u>Figure 1*B*</u>; K**)^4, 5^, thereby challenging the classical dichotomy between clear synaptic vesicles and monoaminergic dense-core vesicles^1, 2^. By segregating histamine and GABA into distinct intravesicular domains, the core–rim architecture represents a previously unrecognized mode of dual-transmitter organization. The histamine core was validated by comprehensive positive and negative controls. Positive controls using established histamine-containing gastric enterochromaffin-like (ECL) cells confirmed robust HA2 immunoreactivity^6–9^. Additional specificity controls in gastric tissue, where histamine- and GABA-positive cell populations are anatomically distinct, further confirmed the specificity of HA2 **(<u>Supplementary Figure 1</u>).** Negative controls—including isotype controls, antigen preabsorption and primary-antibody omission—abolished the specific diaminobenzidine (DAB) reaction. Because amino-acid transmitters do not form dense cores under these fixation conditions^1^, this polarized core–rim architecture represents a previously unrecognized structural principle for histamine/GABA co-packaging.

**Figure 1.**
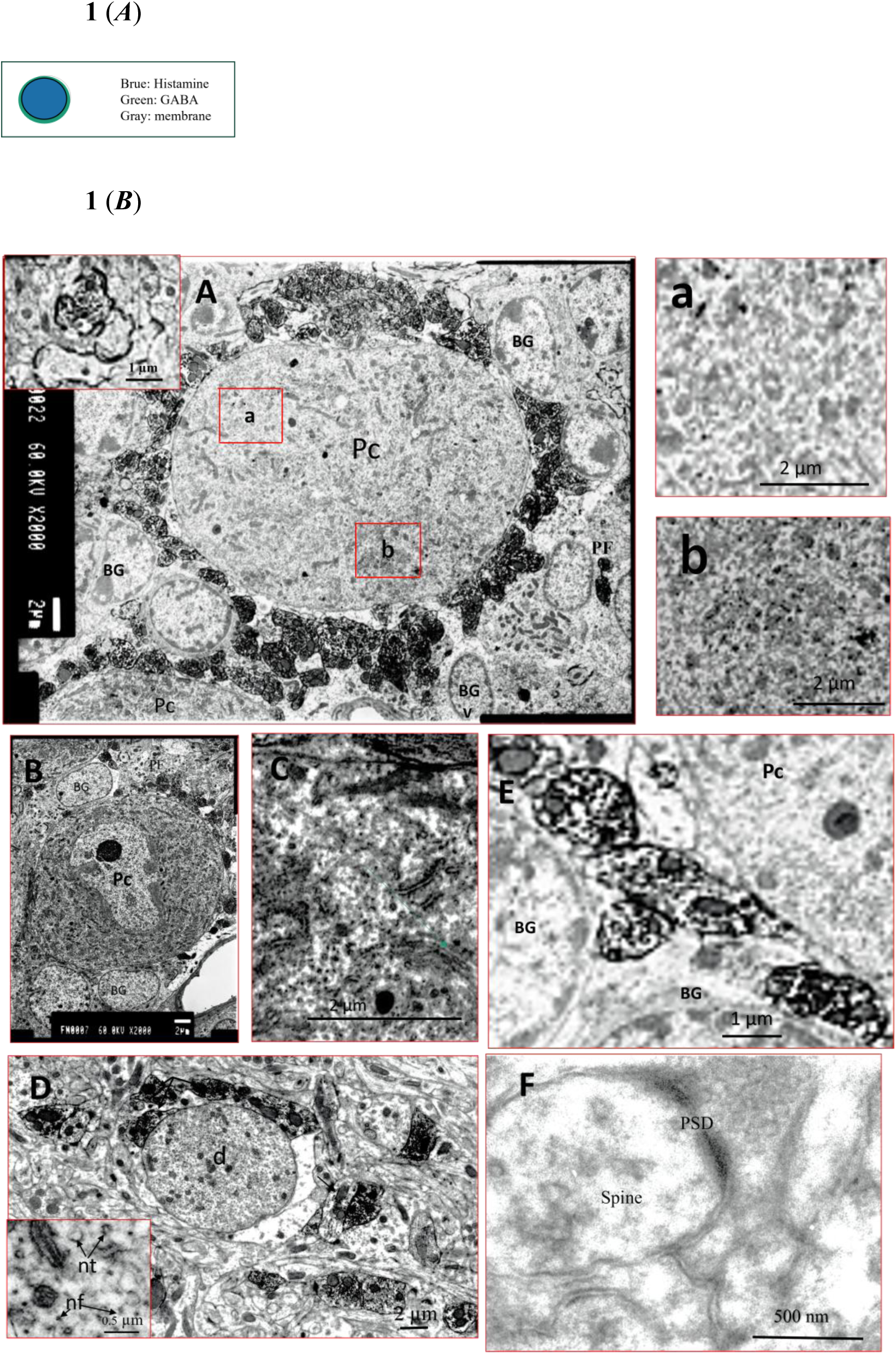
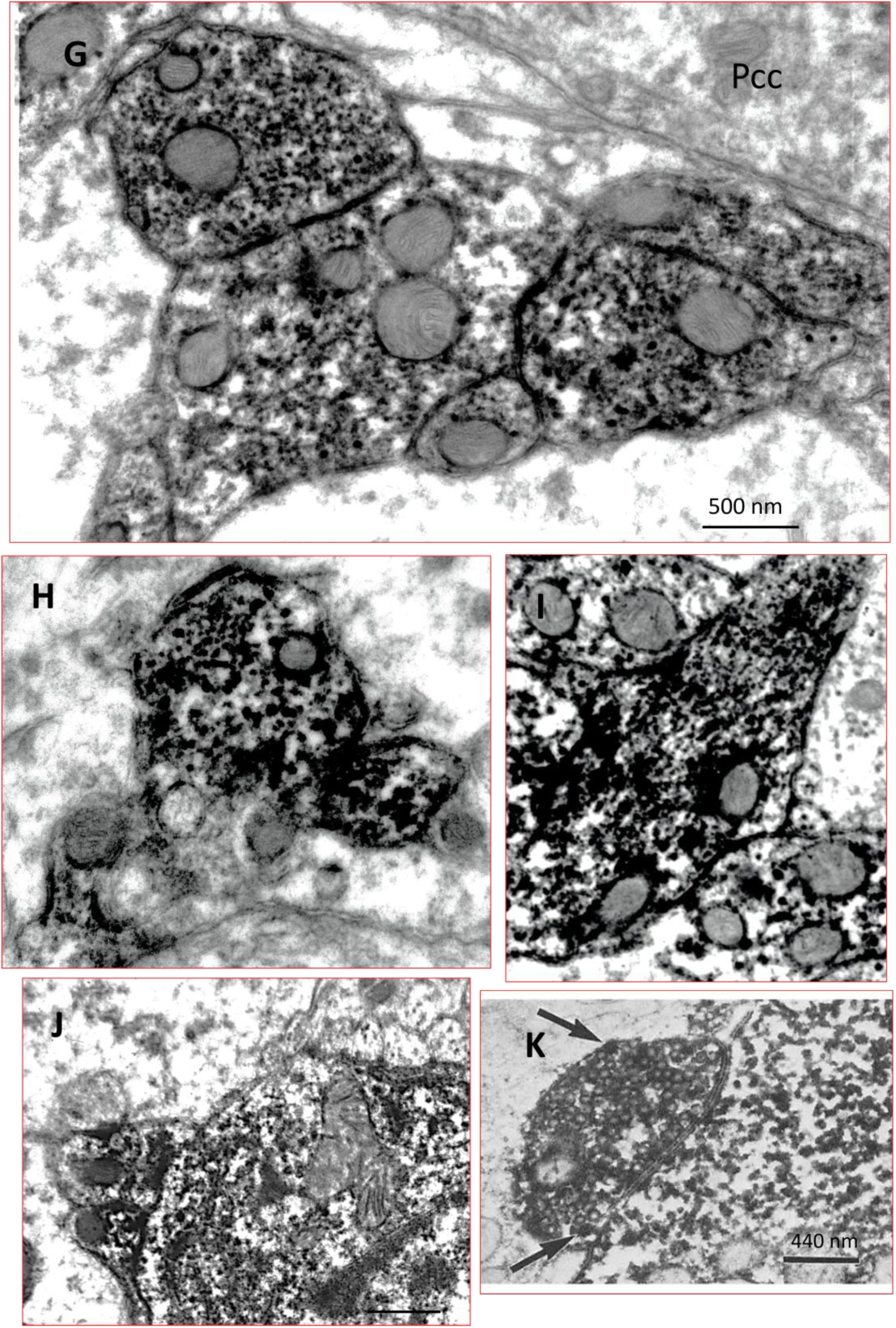
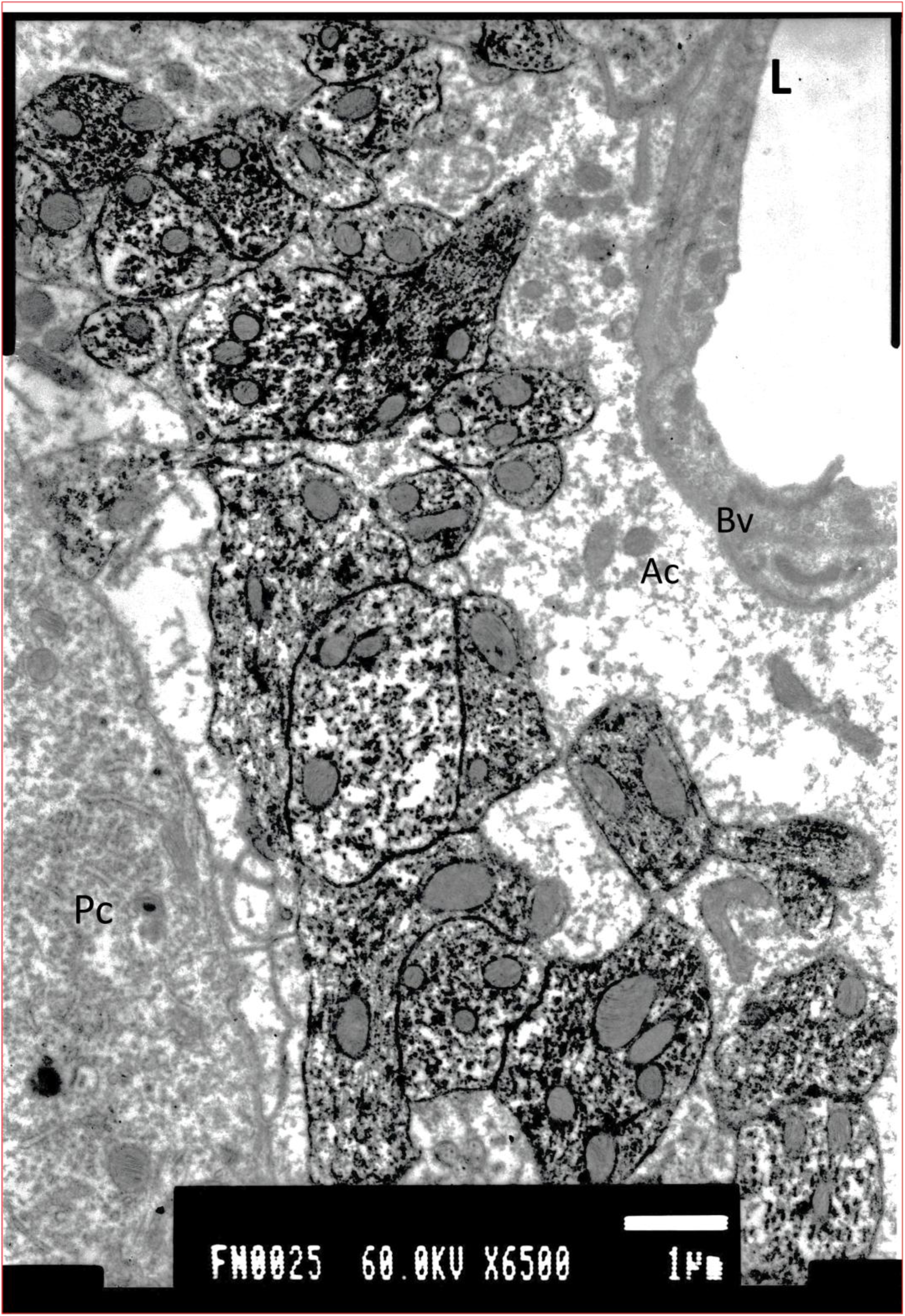
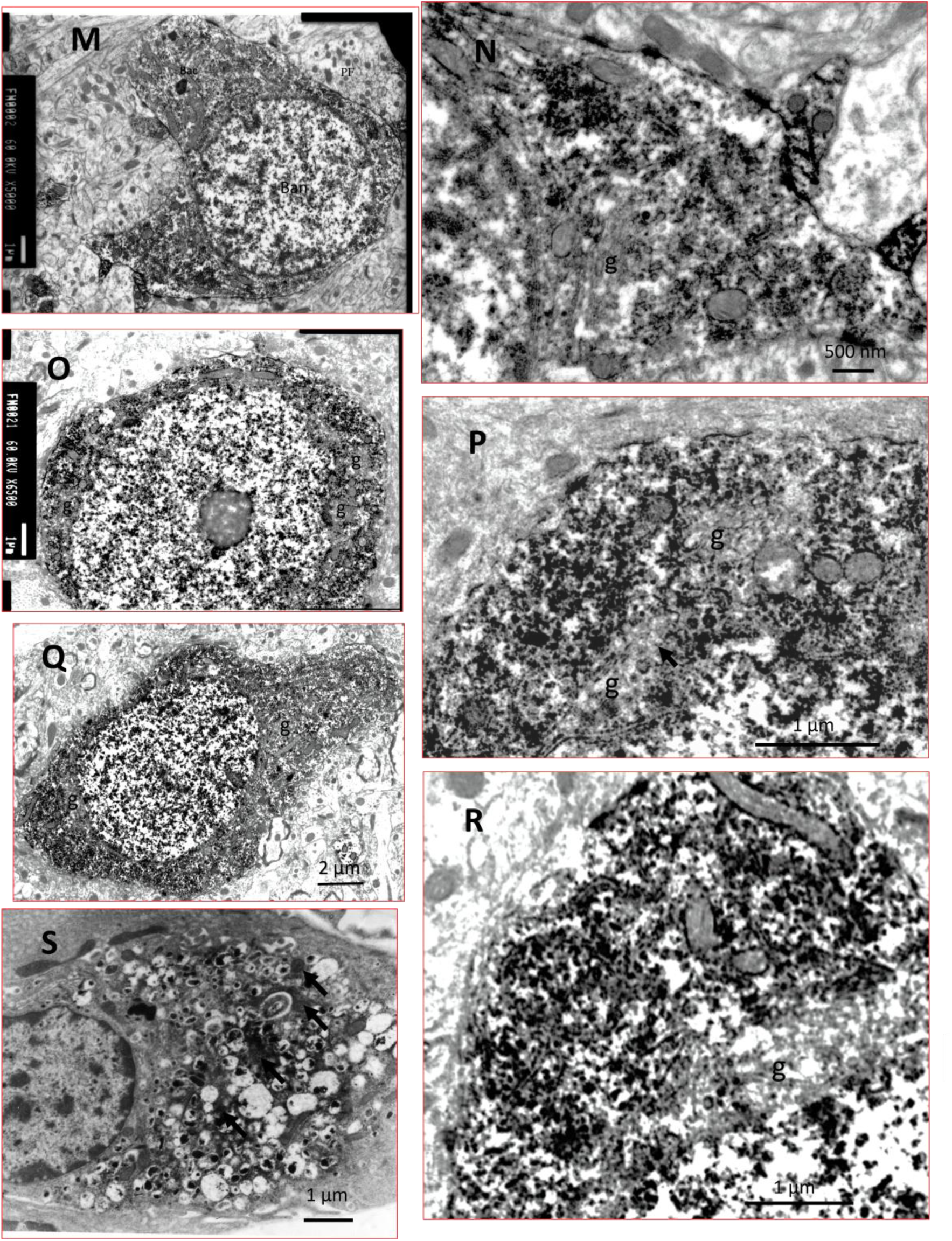
**(A) Schematic representation** of the core–rim vesicular organization. Schematic model illustrating the nanoscale spatial segregation of transmitters within a single synaptic vesicle. Histamine is concentrated within the electron-dense inner core, whereas GABA occupies the membrane-proximal outer rim. **(B) Electron microscopy** reveals a polarized core–rim vesicular architecture. Low-magnification views of Purkinje-related ultrastructure (**A–F**), high-magnification views of core–rim vesicles (**G–K**), cell-type-specific inhibitory profiles (**L–R**), and an endocrine reference (**S)** establish the ultrastructural organization of histamine- and GABA-containing vesicles. Histamine formed a dense central core, whereas GABA was restricted to a thin peripheral rim. Apart from this core–rim organization, vesicular and synaptic morphology was indistinguishable from that of classical GABAergic terminals^5. Radial analysis showed preferential histamine localization within the inner 30-50% of the vesicle radius, and quantitative analysis confirmed predominant core localization, with 2,195 core-associated versus 108 peripheral DAB signals across 23 boutons (95.1% core-associated; Golgi regions excluded) **Purkinje cell layer** (**A–F**). Purkinje cell bodies and dendrites showed weak labeling, whereas ribosomes and short rough endoplasmic reticulum segments were strongly immunoreactive. Nerve filaments and microtubules were weakly labeled. Gray type I synapses containing round vesicles were immunonegative (IN), serving as internal controls^35^. **High-magnification electron micrographs (G–K).** Basket cell terminals contained dense-core histamine-positive vesicles surrounded by a thin GABA-immunolabeled rim. Vesicles frequently aligned along microtubules (**I**). Occasional histamine-positive boutons formed asymmetric contacts with IN profiles (**J**). Panel (**K)** shows the classical peripheral GABA rim for comparison. **Medium magnification near Purkinje somata (L).** Basket cell collaterals formed symmetric Gray type II synapses with neighboring boutons but not with Purkinje somata. Vesicles consistently displayed central histamine labeling. **Basket, stellate, and Lugaro cells (M–R).** These neurons showed strong cytoplasmic labeling with clusters of histamine-positive vesicles around immunoreactive regions. Vesicles within the Golgi apparatus and mitochondrial matrix were unlabeled. **Comparative control (S).** Gastric ECL cells exhibited both dense-core and cytoplasmic HA2 labeling, providing an independent morphological reference. **Abbreviations:** Pc, Purkinje cell; Pcd, Purkinje dendrite; BG, Bergmann glia; Ac, astrocyte; g, Golgi region; d, dendrite. Differences in optical density between the vesicle core and peripheral rim were statistically significant (P < 0.001, Wilcoxon rank-sum test). Scale bars: 200 nm **(A–F),** 500 nm **(G–I, N),** 100 nm **(J),** 440 nm **(K),** 1 µm **(L, M, O, P, R, S),** 2 µm **(Q).**

This core–rim architecture extends the paradigm previously established in gastric enterochromaffin-like (ECL) paraneurons using the identical HA2 antibody (**<u>Figure 1*B*</u>; S**). The conservation of this organization across CNS and endocrine systems is consistent with the established storage principle of other monoamines^2^, which condense into dense cores for stable packaging. By positioning a histamine core within a GABAergic rim, this vesicular organization bridges the structural distinction between monoaminergic sequestration and rapid synaptic transmission architectures^1, 2^.

**Quantitative radial analysis**^10^ confirmed marked histamine enrichment within the inner 30–50% of the vesicular radius, with 95.1% of the total signal confined to the core (*n* = 2,195 versus 108 vesicles across 23 boutons; **<u>Figure 1*B*</u>; G–J, L**). The absence of intraluminal membranes suggests that this nanoscale segregation is not maintained by membrane-bounded diffusion barriers, but may instead arise from intrinsic physicochemical mechanisms, including matrix binding and charge-dependent condensation^11, 12^

This geometrical arrangement may provide a structural framework for temporally staggered transmitter release during a single exocytic event. Upon fusion-pore opening, membrane-proximal GABA would be expected to diffuse rapidly, consistent with its role in fast synaptic transmission. Histaminergic signaling, in contrast, is classically associated with slower, diffuse neuromodulatory transmission rather than conventional rapid synaptic signaling^13^. In this context, the condensed histamine core may dissociate more slowly than the membrane-proximal GABA pool, potentially providing a structural basis for the delayed and sustained component of histaminergic signaling (**<u>Figure 5</u>**). Although release kinetics were not directly measured here, this interpretation links the newly identified intravesicular architecture to the established temporal properties of histaminergic neurotransmission.

**The observable range of neuronal histamine** was previously largely confined to pathways originating from the tuberomammillary nucleus (TMN) ^3, 14^. The GA–NaBH₄ method expands this experimentally accessible domain without requiring revision of the established TMN circuitry. The previous view was shaped in part by historical technical limitations: early antibodies generated against fetal HDC lacked the sensitivity required to detect low-level HDC expression in the adult brain and peripheral tissues. Similarly, early histamine mapping efforts were constrained by the use of formaldehyde fixation—a method believed to result in poor histamine retention^14^.

**Antigen–antibody interactions** generally require an epitope of at least the molecular size of a dipeptide or trisaccharide for stable molecular recognition^15, 16^. We developed an epitope-engineering strategy targeting aliphatic primary amines, a pioneering platform that has since been extended to the ultrastructural localization of polyamines^17^. and exogenous drugs^18,19^. Because histamine is too small to serve as a conventional antibody epitope, specificity depends on recognition of the histamine–GA moiety generated within fixed protein complexes through the same chemistry that underlies in situ fixation. NaBH₄ reduction is a critical step in this detection strategy: by converting residual reactive aldehydes to non-reactive alcohols, it substantially suppresses nonspecific background reactivity while preserving the GA-derived histamine epitope, thereby enabling specific visualization of histamine in fixed tissue. The histamine–GA–BSA immunogen generates at least two GA-dependent epitope classes recognized by distinct monoclonal antibodies, HA1/3–5 and HA2^6–9^. Whereas gastric enterochromaffin-like (ECL) cells express both epitope classes^6–9^, neural tissue appears to generate only the HA2-type epitope^20^. The proposed chemical nature of this HA2 epitope is illustrated in **<u>Supplementary Figure 2</u>**.

**Retaining highly labile small-molecule histamine** necessitates robust GA fixation, which intrinsically generates a dense cross-linking network that severely restricts tissue permeability. Our pre-embedding DAB platform overcomes this constraint sufficiently to permit antibody access to fixation-preserved vesicular compartments, where localized immunoreactivity is converted into electron-dense DAB reaction product that can be resolved at ultrastructural resolution. In contrast, conventional post-embedding immunogold labeling is poorly suited to this system. The limited penetration of antibody–gold complexes into heavily cross-linked tissue, together with the finite size of the immunogold probe and associated antibody complex (approximately 15–25 nm), imposes substantial spatial uncertainty relative to the dimensions of individual synaptic vesicles and therefore cannot reliably resolve the core–rim organization described here^21, 22^. Accordingly, immunogold-based double labeling of histamine and GABA within the same synaptic vesicles is not technically feasible under these conditions^21, 22^..

**The high signal-to-noise performance** of the GA–NaBH₄ method enabled ultrastructural visualization of transmitter-containing vesicles within neuronal somata. Notably, even the most extensively characterized GABAergic vesicular systems have been examined predominantly in presynaptic terminal boutons, with vesicles within neuronal somata not previously resolved at comparable ultrastructural resolution. The enhanced signal-to-noise performance of our method enabled clear visualization of histamine-immunopositive (IP) synaptic vesicles within the cell bodies of Purkinje, basket, stellate, and Lugaro cells, as well as along their axons and within presynaptic terminals (**<u>Figure 1*B*</u>**; **A–R**). The continuous occurrence of histamine-IP vesicles across somata, axons, and presynaptic terminals provides morphological evidence consistent with their somatic origin and subsequent axonal transport to distal synaptic sites. Thus, the GA–NaBH₄ method provides sufficient sensitivity and contrast to resolve vesicular organization within the cytoplasm of neuronal somata and along axonal trajectories, extending ultrastructural analysis beyond the conventional terminal-bouton compartment.

**To determine** whether these molecular signatures are reflected in the localized end-product histamine, we examined the spatial distributions of GABA and histamine by high-resolution light microscopy. Immunohistochemical analysis of adult rat cerebellar sections revealed a strikingly similar spatial pattern for the two transmitters (**<u>Figure 2 *A*</u> and *B***). Within the cerebellar cortex, Purkinje cell (PC) somata exhibited faint but detectable immunoreactivity for both GABA and histamine, with the surrounding neuropil showing stronger labeling (**<u>Figure 2*B*</u>**; **G, H, g, h**). In the deep cerebellar nuclei (DCN), terminals immunoreactive for both transmitters formed a thin pericellular outline around the somata of large projection neurons (**<u>Figure 2*B*</u>**; **I, i**). Thus, both GABA and histamine displayed a consistent spatial configuration characterized by relatively weak somatic labeling and a more prominent peripheral signal, extending from neuronal somata to their surrounding terminal fields.

**Figure 2.**
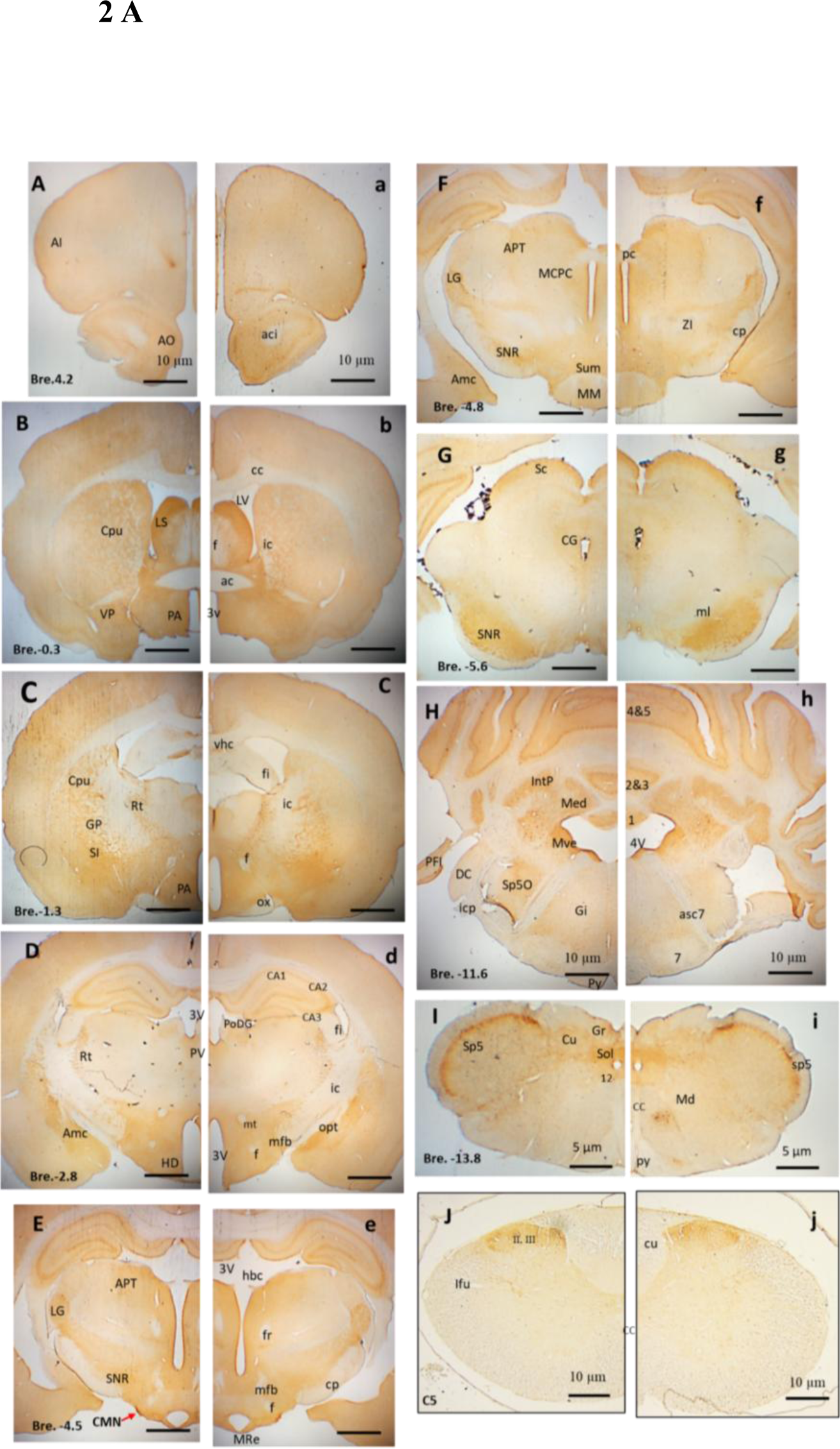
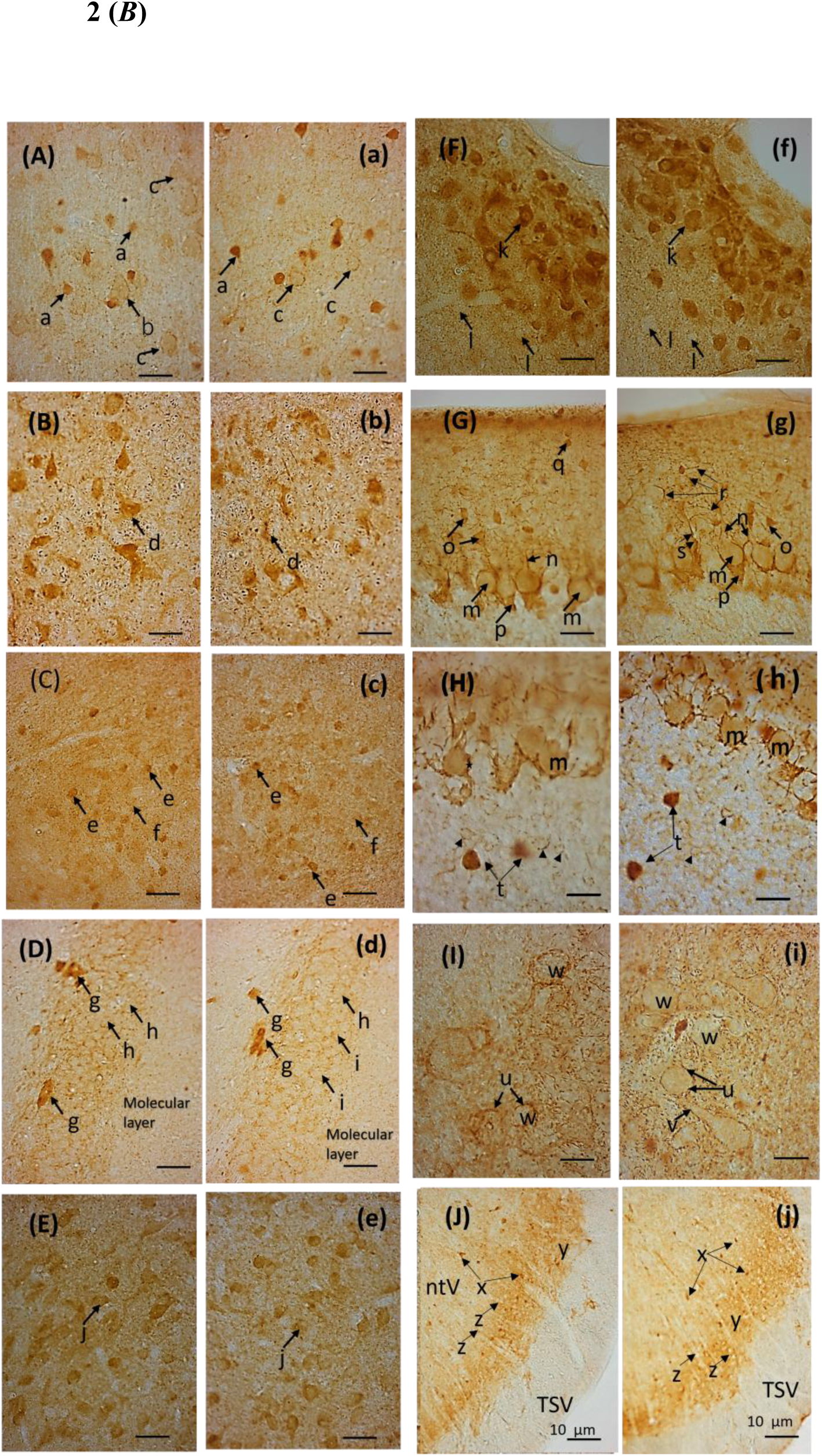
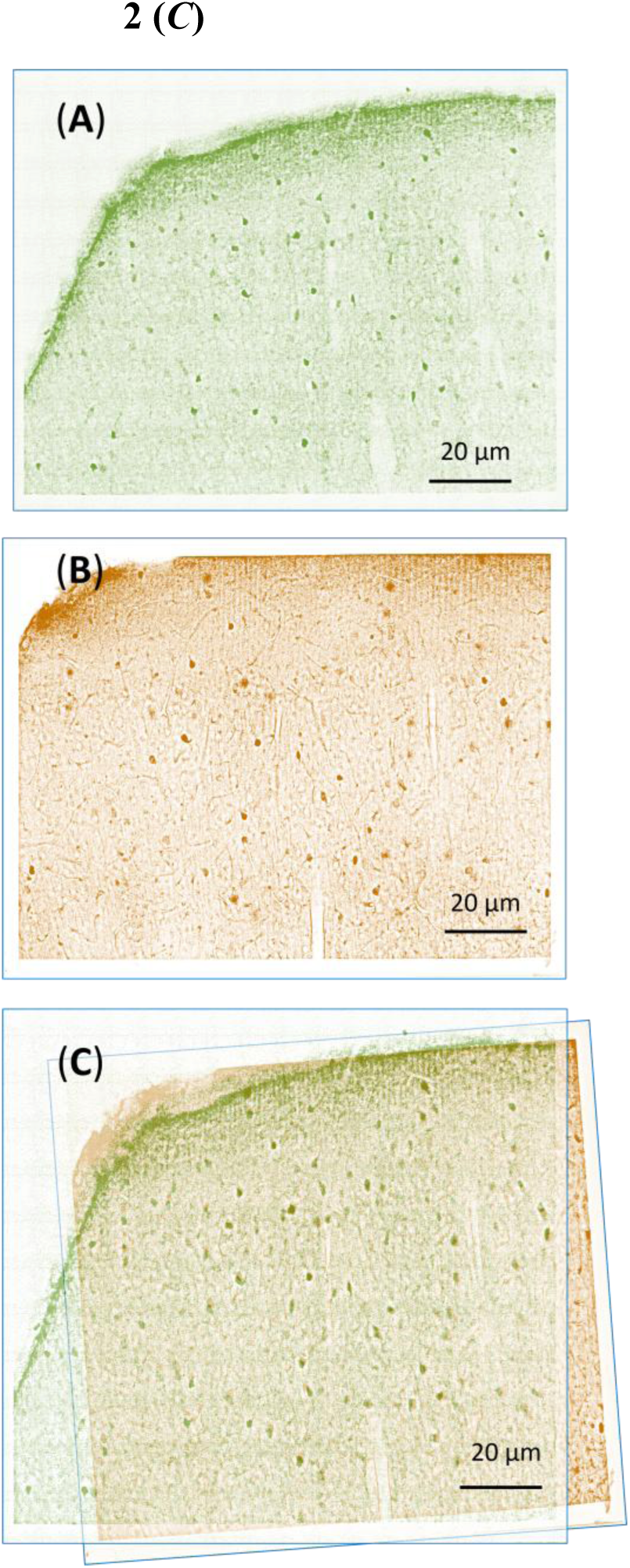
Core–rim organization and histamine/GABA colocalization in inhibitory neurons. This figure is organized into three hierarchical panels: whole-brain distribution (***A***), (***B***) high-magnification cellular profiles, and (***C***) histamine–GABA colocalization. Together, these panels provide the light-microscopic foundation for the polarized core–rim architecture revealed by electron microscopy. **(A) Distribution of histamine (A–J) and GABA (a–j) immunoreactivity in serial frontal sections processed under identical conditions.** Histamine-positive elements exhibit a spatial organization that closely parallels the GABAergic system with anatomical localization referenced to the standard Paxinos and Watson stereotaxic atlas^36^. Panels **A–J** and **a–j** are corresponded to frontal sections at the indicated Bregma levels. **Abbreviations**: AI, agranular insular cortex; Amc, amygdaloid nu complex; AO, anterior olfactory nu; APT, ac, anterior commissure; anterior pretectal nu; aci, anterior commissure, intrabulbar part; asc7, ascending fibers of the facial nerve; CA1-3, fields CA1-3 of Ammon’s horn; CMN, caudate magnocellular nu; Cpu, caudate putamen; CG, central gray; Cu, cuneate nu; cc, corpus callosum; cp, cerebral peduncle; CC, central canal; cu, cuneate fasciculus; DC, dorsal cochlear nu, oral; f, fornix; fi, fimbria hippocampus; fr, fasciculus retroflexus; Gi, giantcellular reticular nu; GP, globus pallidus; Gr, gracile nu; HD, hypothalamic nu; hbc, habenular commissure; Ifu, lateral funiculus spinal cord; IntP, interposed cerebellar nu; ic, internal capsule; icp, inferior cerebellar peduncle; LG, lateral geniculate nu; LS, lateral septal nu; LV, lateral ventricle; MCPC, magnocellular nu of the posterior commissure; Med, medial cerebellar nu; Md, medullary reticular nu; MM, mammillary nu; Mre, mammillary recess; MVe, medial vestibular nu; mfb, medial forebrain bundle; ml, medial lemniscus; mt, mammillothalamic tract; opt, optic tract; ox, optic chiasm; PA, preoptic area; PoDG, polymorph layer of the dentate gyrus; PV, paraventricular nu; PFI, paraflocculus; pc, posterior commissure; py, pyramidal tract; Rt, reticular thal nu; Sc, superior colliculus; SI, substantia innominate; SNR, substantia nigra reticular; Sol, nu solitary tract; Sp5, spinal trigeminal nu; Sp5O, spinal trigeminal nu; Sum, supramammillary nu; sp5, spinal trigeminal tract; VP, ventral pallidum; 3V, 3rd ventricle; 4V, 4th ventricle; vhc, ventral hip commissure; ZI, zona incerta; 7, facial nu; 12, hypoglossal nu; 1,2,3,4,5, cerebellar lobules; II, III spinal cord layers. Scale bars: 10 μm (**A-H**, **J**; **a-h, j)**, 5 µm (**I**; **i**) **(B)** Representative histamine (A–J) and GABA (a–j) immunoreactivity at higher magnification. **(A, a)** Cerebral Cortex: Layer III–IV. HA/GABA-positive non-pyramidal cells (a) are observed. Intense punctate terminals outline the perikarya of large immunonegative (IN) pyramidal cells (b) and round cells (c). **(B, b)** Ventral Pallidum: Large pleomorphic neurons (d) exhibit robust immunoreactivity. Note the presence of numerous IP axial cylinders within myelinated fibers. **(C, c)** Amygdala: Medium-sized amygdaloid neurons show moderate (e) to weak (f) immunoreactivity. **(D, d)** Hippocampus: Highly IP polymorphic cells (g) are prominent. IN granule cells (h) are densely delineated by IP punctate dots (i). **(E, e)** Lateral Septal Nucleus: Moderate staining in medium-sized neurons (j) and their associated neuropil. **(F, f)** Caudal Magnocellular Nucleus (CMN): These neurons display the most intense immunoreactivity (k) in the brain. IP dots outline neighboring IN neurons (l) within the neuropil. **(G, g)** Cerebellar Paraflocculus (Molecular Layer): Purkinje cell bodies (m) and dendrites (n) are weakly labeled, whereas basket cells (o) and stellate cells (q) are strongly immunopositive. Characteristic pinceau formations (p), varicose puncta (r), and basket cell terminals (s) are evident. **(H, h)** Cerebellar Granular Layer: Strong IP staining in Golgi cells (t) and punctate profiles (arrowheads) at the periphery of cerebellar glomeruli. Granule cell clusters remain IN. **(I, i)** Deep Cerebellar Nucleus: IP bouton-like axon terminals (likely Purkinje-derived) punctately surround the somata (u) and proximal dendrites (v) of large IN neurons (w). **(J, j)** Spinal Trigeminal Nucleus: Robust immunoreactivity in cells (x) and neuropil (y) of the substantia gelatinosa. Neighboring cells (z) and the spinal trigeminal tract (TSV) show minimal reactivity. Scale bars: 2.5 µm (**A–I; a–i**); 10 µm (**J, j**) **(C)** Colocalization of histamine and GABA in inhibitory neurons. Free-floating double immunostaining demonstrated complete somatic colocalization of histamine and GABA in neocortical inhibitory neurons, with corresponding dendritic and terminal labeling. Neocortex, Bregma −1.3 mm; Scale bars: 20 μm **(A–C)**

**In the cerebral cortex,** optimized tissue permeabilization enabled high-resolution DAB double staining^23^ and revealed complete cellular overlap between histamine- and GABA-immunopositive neuronal populations. Among clearly identifiable neuronal somata, all histamine-immunopositive somata were also GABA-immunopositive (110 of 110 cells; 100% overlap) (**<u>Figure 2*C*</u>**). Thus, within the clearly identifiable neuronal population, histamine immunoreactivity was completely concordant with the GABAergic phenotype at the light-microscopic level.

**This pattern extended** beyond the central nervous system to peripheral neural-crest–derived structures. In the adrenal medulla, histamine immunoreactivity was detected in chromaffin cells arranged in cord-like clusters (**<u>Figure 3; a, b</u>**), resembling the established cellular organization of GABA-associated adrenal chromaffin cells^24, 25^. Faint residual staining was observed in some control sections even in the absence of the primary antibody; however, the characteristic cellular pattern observed with HA2 was absent under these conditions (**<u>Figure 3; a′, b′</u>**), supporting the specificity of the histamine-associated signal. In sympathetic ganglia, histamine immunoreactivity was localized to neuronal somata and to incoming nerve fibres forming pericellular networks around ganglion cells (**<u>Figure 3; c, d</u>**), closely paralleling the characteristic GABAergic organization described in peripheral ganglia^26^. Together, these findings strongly suggest that the histamine-associated cellular patterns observed in the CNS also exist within peripheral neural circuits.

**Figure 3.**
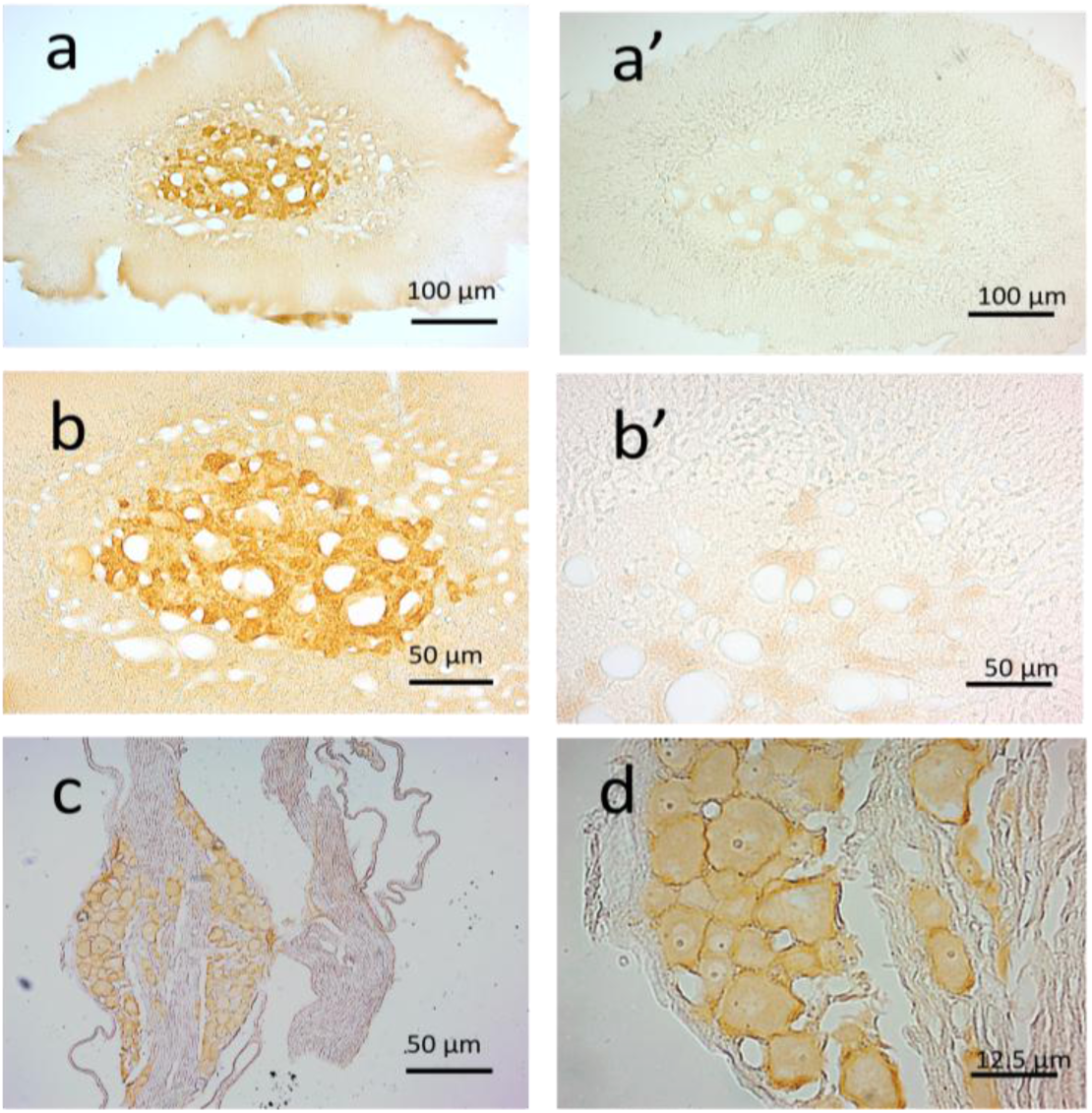
Conservation of the core–rim histamine/GABA organization in endocrine and autonomic systems. **Three components: (a–b)** chromaffin cell histamine storage, **(a′–b′)** HA2 absorption controls, and **(c–d)** sympathetic ganglia varicose fibers—showing conserved histamine/GABA core–rim organization in peripheral systems. **(a, b) Adrenal medulla.** Histamine immunoreactivity exhibits a punctate cytoplasmic pattern in chromaffin cells, consistent with ultrastructural evidence that histamine is stored in VMAT2-dependent dense-core vesicles and with classical studies of monoamine storage in endocrine cells^24, 25^. **(a′, b′) Absorption controls.** Preabsorption with histamine–GA–BSA and omission of the primary antibody both eliminated specific immunostaining. The faint residual coloration occasionally observed in noradrenaline (NA) cells was unaffected by either control, indicating that it reflects intrinsic oxidation products rather than incomplete antibody absorption or antibody-dependent staining. **(c, d) Sympathetic ganglia.** HA2-positive varicose terminals formed a dense pericellular network around principal neurons, whereas neuronal somata showed only weak immunoreactivity. Numerous histamine-positive fibers coursed between ganglionic cell bodies, forming a characteristic pericellular innervation pattern consistent with autonomic cotransmission^26^. Together, these findings demonstrate that histamine/GABA cotransmission extends beyond the central nervous system to autonomic and endocrine tissues. Histamine-positive varicose fibers in sympathetic ganglia and VMAT2-dependent dense-core vesicles in adrenal chromaffin cells indicate a conserved vesicular organization across peripheral effector systems. **Scale bars:** 100 μm (**a, a′, c**), 50 μm (**b, b′**), 12.5 μm (**d**).

**How cytosolic histamine (HA) and GABA** are incorporated into the same vesicle remains unknown. The most straightforward model is that VGAT and VMAT2 mediate vesicular loading of GABA and HA, respectively. Alternatively, VMAT2 may contribute to the transport of both transmitters, as reported for GABA in dopaminergic neurons^27,28^, while organic cation transporters may also contribute to intracellular HA accumulation^29^.Following transporter-mediated uptake, intravesicular physicochemical interactions, such as matrix binding and charge-dependent condensation, could drive the preferential localization of HA to the vesicular core and GABA to the peripheral rim. Thus, the newly identified core–rim architecture may reflect the integration of established vesicular transport mechanisms with subsequent intraluminal physicochemical partitioning. **Analysis of the Macosko cerebellar dataset**^30^ revealed cell-type-specific *Hdc* transcript detection across the four major cerebellar cell classes. Purkinje cells showed the highest *Hdc* signal, with a mean expression of 0.912 and *Hdc* detected in 15.51% of cells (130/838). Molecular layer interneurons (MLIs) expressing *Megf11* (multiple EGF-like domains protein 11) showed lower but detectable *Hdc* expression (mean, 0.149; 1.92% positive), whereas *Cdh22*-expressing MLIs and granule cells showed little to no detectable *Hdc* (0.22% and 0.008% positive, respectively) (**<u>Figure 4</u>**). Thus, among the cerebellar populations examined, Purkinje cells showed the clearest transcriptional evidence for *Hdc* expression.

**Figure 4.**
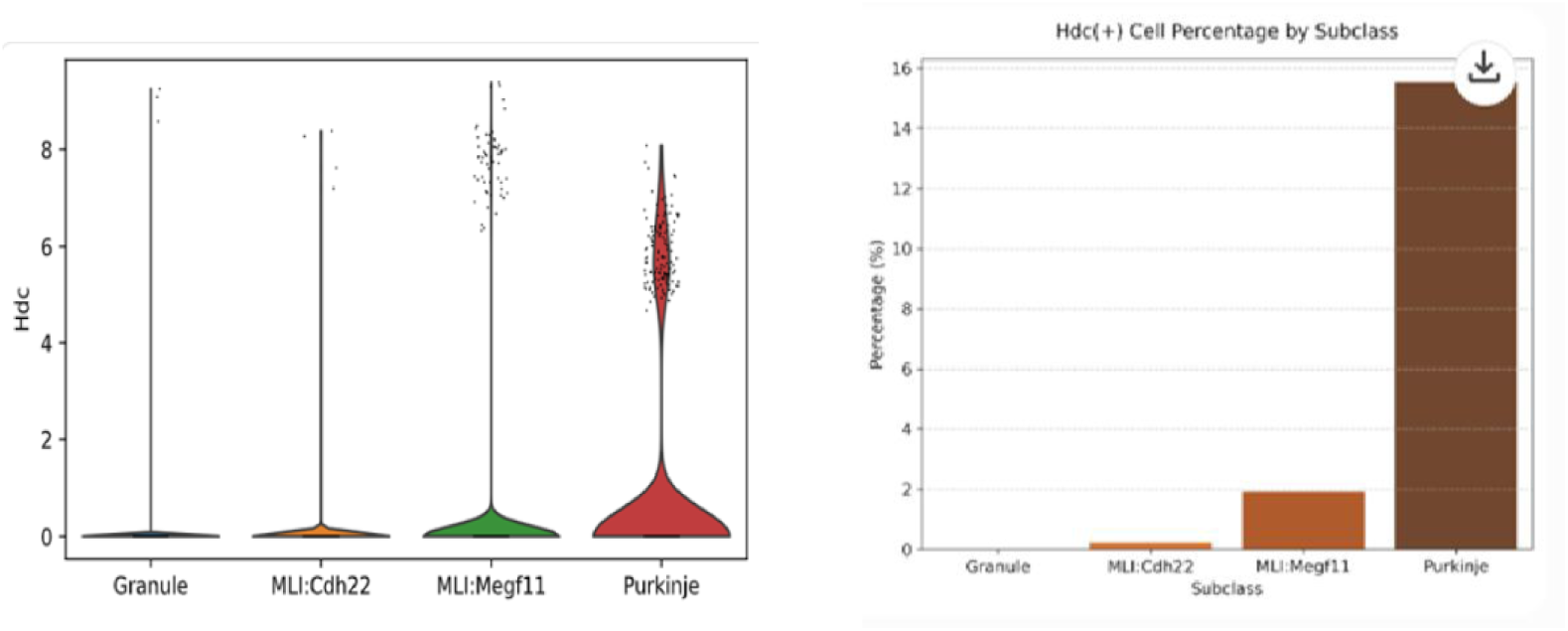
Analysis of *Hdc* expression and cellular proportion across cerebellar subclasses. Two components are shown: (Left) single-cell *Hdc* expression across cerebellar subclasses, and (Right) the proportion of *Hdc*-positive cells. Together, these panels indicate that Purkinje cells contain a distinct subset of *Hdc*-expressing neurons. **(Left)** Violin plot showing the distribution of *Hdc* expression levels in individual cells across four cerebellar subclasses: Granule, Molecular level Interneurons (MLIs expressing cadherin-22 (Cdh22), MLI expressing multiple EGF-like domains protein 11 (Megf11), and Purkinje. Each dot represents a single cell. **(Right)** Bar chart illustrating the percentage of *Hdc*-positive (*Hdc* (+)) cells within each subclass. The Purkinje cell subclass exhibits the highest proportion of *Hdc* (+) cells compared to the other groups.

**The Allen (Macosko)** dataset revealed low-frequency yet distinct *Hdc* expression (15.5%) in Purkinje cells, providing independent molecular evidence that histamine-synthesizing capacity exists in neurons outside the tuberomammillary nucleus (TMN) (**<u>Figure 4</u>**). Crucially, while the present study detects histamine itself, the Allen dataset measures *Hdc* transcripts; levels of transcripts and histamine may diverge due to factors such as translation efficiency, cofactor availability, or vesicle dynamics^31–34^. The approach utilizing the GA–NaBH₄ method offers complementary evidence enabling histamine detection beyond the traditional confines of the TMN, revealing a spatial organization not fully captured by current transcriptome models.

**These findings** reveal a previously unrecognized vesicular structural principle that may help explain the established physiological properties of GABA and histamine. The membrane-proximal localization of GABA within the vesicular rim is consistent with its rapid, phasic inhibitory action, whereas the condensation of histamine within the vesicular core may contribute to its slower mobilization and release, consistent with the slower and sustained neuromodulatory signaling characteristic of histamine, including volume transmission. Our structural findings do not establish the mechanisms of volume transmission itself; rather, they identify an intravesicular organization that may contribute to the distinct release properties underlying this mode of signaling. Thus, the newly identified nanoscale organization provides a structural basis for understanding the long-standing functional disparity between GABAergic and histaminergic signaling, linking transmitter-specific signaling kinetics to a previously unrecognized intravesicular structural principle **(<u>Figure 5</u>).**

**Figure 5.**
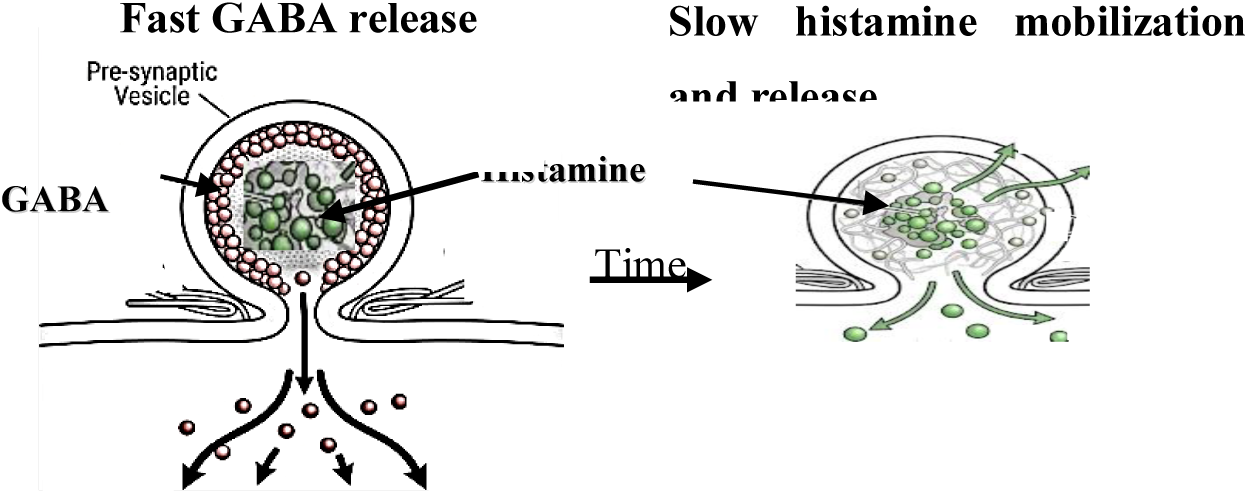
Proposed model of temporally distinct transmitter mobilization from a core–rim vesicle. Schematic working model illustrating how the nanoscale core–rim organization of histamine and GABA within a single synaptic vesicle may support temporally distinct transmitter signaling. Membrane-proximal GABA in the peripheral rim is proposed to become rapidly available upon vesicle fusion, whereas histamine condensed within the vesicular core may require slower mobilization for release. The model links the observed spatial organization of GABA and histamine to their established differences in signaling dynamics. This model is conceptual and does not represent directly measured release kinetics.

**In summary,** this study reveals a previously unrecognized “core–rim” structural polarity within individual synaptic vesicles, characterized by a histamine core and a GABA rim. This organizational pattern appears to be conserved from the central nervous system to the periphery, challenging the traditional dichotomy between clear synaptic vesicles and dense-core vesicles. This nanoscale organization may provide a structural basis for, and help explain, the well-established physiological differences between histamine and GABA signaling, including the slower, sustained action of histamine compared with the rapid inhibitory action of GABA.

Notably, transcriptomic and histochemical signals did not show a simple quantitative correspondence: Purkinje cells, which showed relatively high *Hdc* expression in the Allen dataset, exhibited weak histamine immunoreactivity, whereas molecular-layer interneurons, despite extremely low *Hdc* expression, showed strong histamine immunoreactivity. The biological basis of this apparent discrepancy remains an important physiological question for the next generation of researchers.

## Abbreviations

IHC: Immunohistochemistry
GABA: γ-aminobutyric acid
NaBH₄: sodium borohydride
HDC: histidine decarboxylase
HA2: glutaraldehyde-dependent anti-histamine monoclonal antibody (mAb)
HA05: anti-histamine monoclonal antibody (mAb)
GA: glutaraldehyde
TMN: tuberomammillary nucleus
CNS: central nervous system
ECL: enterochromaffin-like cell
EM: electron microscopy
LM: light microscopy
DAB: 3,3′-diaminobenzidine
IP: immunopositive
IN: immunonegative
VGAT: vesicular GABA transporter
VMAT: vesicular monoamine transporter
*Hdc*: histidine decarboxylase transcripts

## Supplement Data

**Figure 1.**
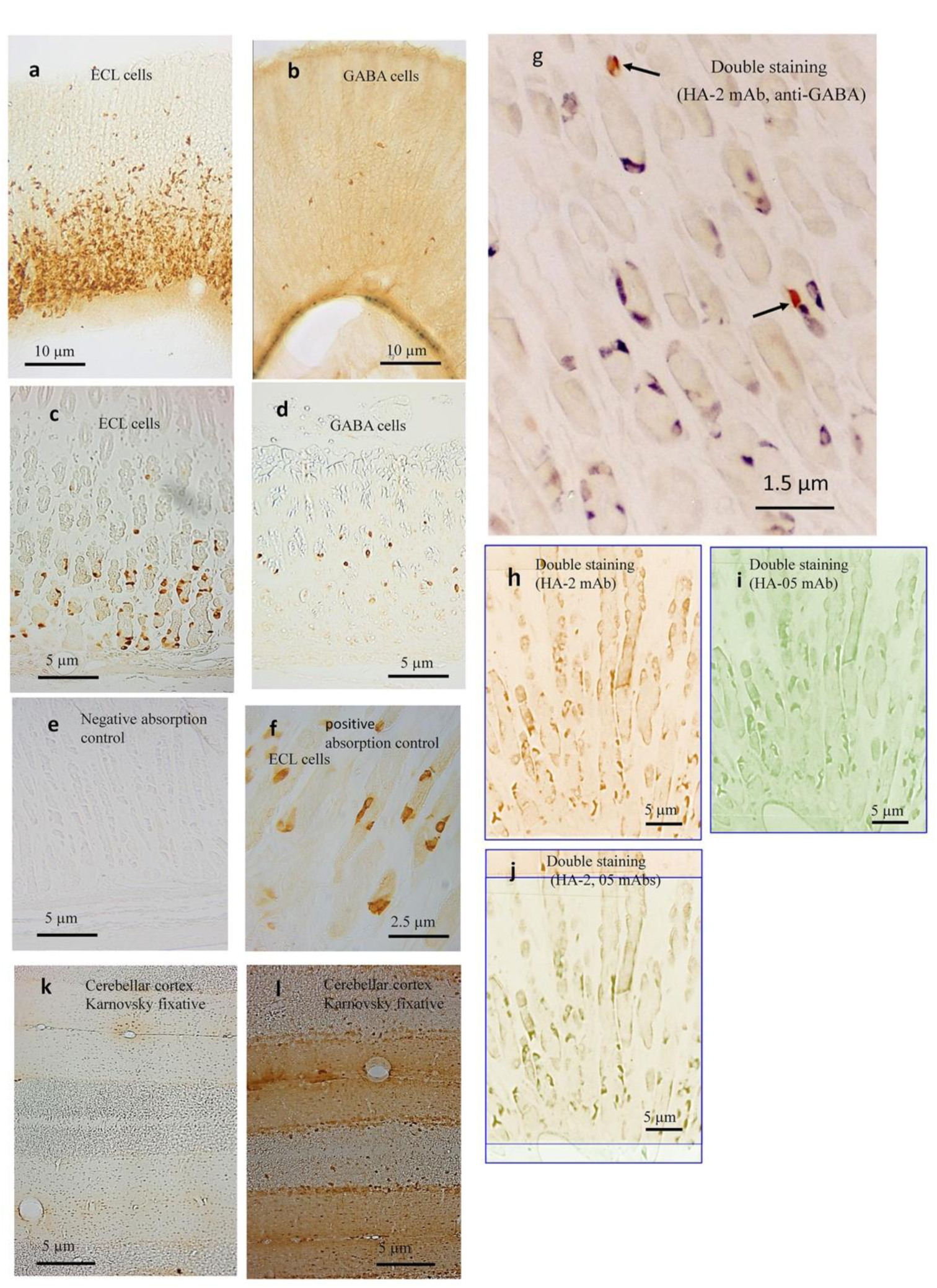
Verification of HA2 antibody specificity. Because histamine and GABA exhibited similar staining patterns in the brain, antibody specificity was verified in gastric fundus tissue, where the two cell populations are clearly distinct. Absorption controls, dual labeling, independent antibody validation, and fixative dependency confirmed the specificity of HA2. **(a)** HA2 selectively labels enterochromaffin-like (ECL) cells clustered in the lower gastric glands. **(b)** GABA mAb labels a sparse, morphologically distinct cell population with thin apical processes. **(c, d)** High-magnification paraffin sections demonstrate the distinct morphologies of histamine-positive ECL cells **(c)** and GABA-positive cells **(d)**. **(e, f)** Preabsorption with histamine–GA–BSA abolishes HA2 immunoreactivity^6,7,8,9,20^, whereas GABA–GA–BSA has no effect. **(g)** HA2 and GABA mAbs label non-overlapping cell populations. **(h–j)** Independent validation using HA2 (**h**) and HA05 (**i**) demonstrates identical histamine labeling patterns^6, 20^. **(k, l)** Karnovsky fixation^37^ abolished HA2 labeling by altering the GA-dependent histamine epitope, whereas GABA immunoreactivity is preserved. **Scale bars:** 10 μm **(a, b);** 5 μm **(c–e, h–j, k, l);** 2.5 μm **(f);** 1.5 μm **(g).**

**Figure 2.**
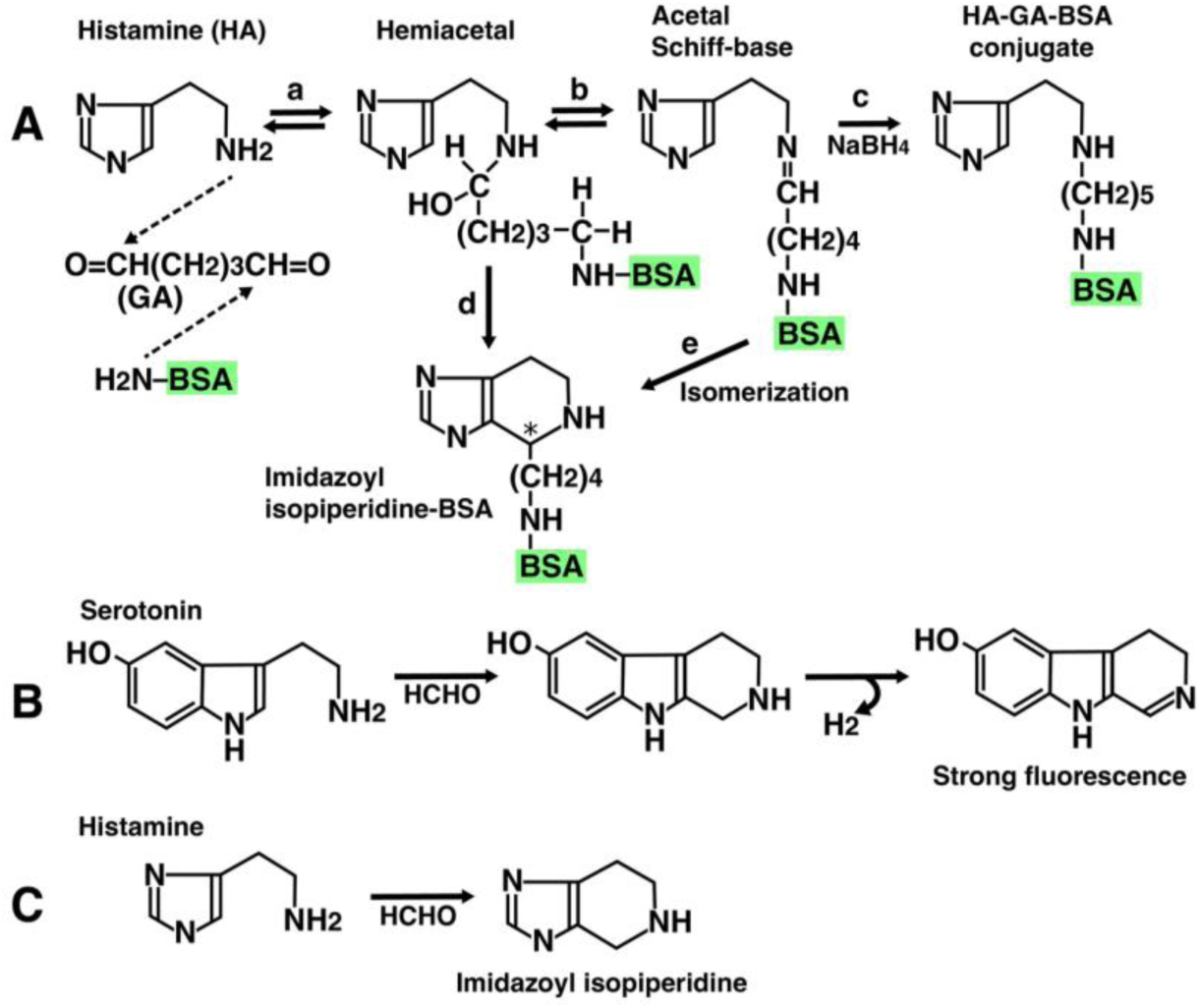
Proposed chemical structures of the glutaraldehyde-dependent histamine epitope. While all five monoclonal antibodies (HA1–HA5) recognize histamine in gastric ECL cells^6,7,8,9^, only HA2 detects histamine in the brain^20^. This difference may stem from the fact that the histamine-GA-BSA antigen may form at least two distinct epitope structures (histamine-GA-BSA or imidazoyl isopiperidine-BSA). The chemical structures of these hypotheses are presented below. (A) Histamine reacts with the aldehyde group at one end of glutaraldehyde to form a hemiacetal, and the reaction proceeds in two directions. In one pathway, a Schiff base is formed, and a histamine-protein complex stabilized with NaBH4, as described in Methods. In the other pathway, the hemiacetal and histamine itself condense to produce an imidazolyl isopiperidine derivative. Furthermore, the Schiff base isomerizes to imidazolyl isopiperidine. This cyclic structure may preferentially form under intracellular conditions within nerve cells, as demonstrated by Falck-Hillarp reaction^38^. (B) Serotonin undergoes a Falck-Hillarp-type aldehyde condensation reaction, producing a cyclic piperidine fluorescent product^38^. (C) Histamine similarly reacts with formaldehyde to produce an imidazolyl isopiperidine structure similar to the classical monoamine-aldehyde reaction^39^. Among these possible reaction products, the imidazolyl-isopiperidine derivative may represent the epitope preferentially recognized by HA2, based on the established chemistry of aldehyde-induced condensation of histamine and related monoamines. This principle may be particularly relevant to small molecules containing multiple reactive functional groups, such as histamine and polyamines^17^, in which fixation and derivatization can generate chemically distinct epitopes and thereby confer clone-specific immunoreactivity.

## Methods and Materials

### Animals and Ethical Statement

Adult male Wistar rats (200–300 g; Kyudo Experimental Animals, Kumamoto, Japan) were housed at 21 ± 1 °C under a 12-h light/dark cycle with food and water available *ad libitum*. All procedures complied with national guidelines and were approved by the Sojo University Ethics Review Committee for Animal Experimentation.

### Tissue Preparation and Fixation

Rats were deeply anesthetized with sodium pentobarbital (60 mg/kg, i.p.) and transcardially perfused with phosphate-buffered saline (50 ml/min, 2 min), followed by freshly prepared 5% glutaraldehyde (GA) in 10 mM phosphate buffer (pH 7.2) for 6 min. Brains were removed, post-fixed overnight at 4 °C in the same fixative, and sectioned at 50 μm using a Microslicer (Dosaka EM, Kyoto, Japan). Alternate sections were processed for histamine (HA) or GABA immunohistochemistry.

### Immunohistochemistry (IHC)

IHC was performed as previously described^6,7,8,9,17,18,19,20^. To eliminate non-specific background from residual reactive aldehydes, all sections were treated with 0.2% NaBH₄ for 10 min.

For histamine (HA) staining, free-floating sections were incubated for 48 h at 4 °C with the monoclonal antibody HA2 (1:500–1:1,000) in 50 mM Tris-HCl-buffered saline (TBS, pH 7.4) containing 0.1% saponin, 0.25% bovine serum albumin (BSA), and 1% normal goat serum. Sodium metabisulfite (0.1% Na₂S₂O₅) was included to stabilize the antigen. Sections were then incubated with HRP-conjugated goat anti-mouse IgG/Fab′ (1:200; MBL, Nagoya, Japan) for 12 h at 4 °C, and immunoreactivity was visualized with diaminobenzidine (DAB) and H₂O₂.

GABA immunostaining was performed using the same protocol with a monoclonal anti-GABA antibody (Funakoshi, Japan). For 5-μm paraffin sections of gastric tissue, protease digestion (Type XXIV bacterial protease) was included to enhance antibody penetration^6,7,8,9^.

### Immunoelectron Microscopy

Following the free-floating DAB reaction, specimens were post-fixed with 1.0% osmium tetroxide in 50 mM cacodylate buffer (pH 7.4) for 1 h. Tissues were dehydrated through a graded ethanol series, cleared in propylene oxide, and embedded in Epon 812 resin. Regions of interest were excised using a 2-mm diameter punch, mounted on Epon blocks, and processed into ultrathin sections. Sections were carbon-coated and examined using a 100CX electron microscope (JEOL, Tokyo, Japan) ^7,8,9,17,19^.

### EM Quantification of Vesicular Core Localization

Tissue samples were processed for pre-embedding DAB immunoperoxidase labeling (HA2), post-fixed with osmium tetroxide, embedded in Epon, and cut into ultrathin sections (60–70 nm) for transmission electron microscopy. Only intact, non-overlapping vesicles were analyzed, with immunonegative asymmetric synapses serving as internal controls. DAB intensity was measured along radial transects and normalized to the vesicle radius (0–100%). The vesicle core was defined as 30–50% of the radius, and the peripheral region as 50–100%. Across 23 boutons, a total of 2,195 core and 108 peripheral DAB signals were quantified, yielding a core fraction of 95.1%. Statistical significance was assessed by permutation analysis (10,000 iterations) using randomized radial distributions as the null model, and bootstrap resampling (10,000 iterations) was performed to estimate confidence intervals.

### Double-Staining Immunohistochemistry^23^

Co-localization was examined by sequential immunostaining as previously described. Histamine was first visualized with 4-chloro-1-naphthol (0.02% in TBS containing 0.005% H₂O₂ and 2% ethanol). After digital imaging, bound antibodies were stripped with 0.1 M glycine-HCl (pH 2.2; four 30-min incubations), followed by ethanol decolorization. Sections were then blocked overnight and reprocessed for GABA immunohistochemistry using the DAB method.

### Specificity and Control Experiments

Specificity was verified by (i) omission of the primary antibody, (ii) incubation with an isotype-matched irrelevant monoclonal antibody (anti-penicillin IgG1, 30–80 ng/ml), and (iii) preabsorption of HA2 with histamine–GA–BSA conjugate (5 μg/ml). All controls abolished or excluded specific HA2 immunoreactivity, confirming its high specificity for GA-fixed histamine.

### Single-nucleus RNA sequencing (snRNA-seq) Analysis of Cerebellar Subclasses

To investigate the expression profile of *Hdc* across distinct cerebellar cell populations, a single-nucleus RNA sequencing (snRNA-seq) dataset generated by the Macosko laboratory was analyzed^30^. The dataset, obtained through the Allen Brain Cell (ABC) Atlas, contains transcriptomic profiles of adult mouse brain nuclei, classified according to the mouse brain taxonomy.

Downstream bioinformatic analysis was performed using Scanpy (v1.11.5). For cell type identification, the high-level annotation metadata provided by the original repository was utilized, focusing on four primary cerebellar subclasses: Granule cells, Molecular Layer Interneurons expressing Cdh22 (MLI:Cdh22), Molecular Layer Interneurons expressing Megf11 (MLI:Megf11), and Purkinje cells.

Gene expression distributions across individual cells in each subclass were visualized using violin plots. The normalized expression values [log2(CPM + 1)] were used to plot the density and distribution for each cell type. To calculate the proportion of *Hdc*-positive [*Hdc*(+)] cells within each subclass, cells with an expression value strictly greater than zero for the *Hdc* transcript were defined as *Hdc*(+) cells. The percentage of these positive cells relative to the total number of cells within each respective subclass was calculated and visualized using a bar chart.

## Ethics statement

Ethics approval for this study was obtained from the Sojo University Animal Experiment Ethics Committee.

## Authors contributions

F.K. designed the experiment. F.K. performed the experiments. F. 1. K. analyzed the data. F. K. drew the photographs and figures. F.K. wrote the manuscript.

## Acknowledgements

I would like to express my deep gratitude to my wife, the late Chiemi Fujiwara, who supported me both materially and emotionally. I would also like to express my deep gratitude to the late Takashi Suemasu, who helped me take the electron microscope photographs, to Professor G. Bay, K. Maeda, M. Matsumoto, and C. Tamura, who helped me with my research, to Professor S. Matsufuji for valuable discussion, to Dr. Yu Liu, who helped me with the Allen analysis, to Y. Shirahama, who helped me with my computer, and to H. Fujiwara, M. Tateishi, and K. Tateishi, who provided their kind support. I am truly grateful.

